# Ghrelin proteolysis by insulin-degrading enzyme

**DOI:** 10.1101/2021.06.10.447714

**Authors:** David D. Bocach, Kierstin L. Jones, Jonathan M. Bell, Qiuchen Zheng, Noel D. Lazo, Jillian E. Smith-Carpenter, Benjamin J. Alper

## Abstract

Here we report proteolysis of synthetic acylated human ghrelin by recombinant human insulin-degrading enzyme (IDE). Kinetic parameters and sites of proteolytic cleavage were evaluated. Ghrelin proteolysis by IDE was inhibited by ethylenediaminetetraacetate (EDTA), a metal chelating agent. Ghrelin proteolysis appears at least somewhat specific to M16 family proteases such as IDE, as the M13 protease neprilysin (NEP) did not exhibit ghrelin proteolysis in this study. A quenched fluorogenic peptide substrate comprising the primary sites of IDE-mediated ghrelin proteolysis (Mca-QRVQQRKESKK(Dnp)-OH; Mca: 7-methoxycoumarin-3-carboxylic acid; Dnp: 2,4-dinitrophenyl) was developed and used to evaluate enzyme specificity and kinetic parameters of proteolysis. Like acyl ghrelin, Mca-QRVQQRKESKK(Dnp)-OH was efficiently cleaved by IDE central to the target sequence. We anticipate that this quenched fluorogenic peptide substrate will be of value to future studies of ghrelin proteolysis by IDE and potentially other peptidases.

## Introduction

Ghrelin is an octanoylated 28 amino acid enteroendocrine hormone involved in mammalian energy homeostasis [Kojima et al. 1999, Tschop et al. 2000, Nakazato et al. 2001, Varela et al. 2011, Müller et al. 2015 and others]. Sartorially named for its role as a growth hormone release (GHREL) factor [Kojima et al. 1999], the peptide is frequently coined a “hunger hormone,” as it promotes feeding and control of feeding patterns in humans and rodents [Nakazato et al. 2001, Wren et al. 2001., Asakawa et al. 2001, Cummings et al. 2001, Tschop et al. 2001, Drazen et al. 2006]. As first characterized in rats, ghrelin derives from a 0.62 kb mRNA transcript [Kojima et al. 1999]. In both humans and rats, the ghrelin gene product yields a 117 amino acid preproghrelin precursor protein, which is subject to extensive post-translational processing by endoproteolytic cleavage and acylation [Kojima et al. 1999].

Post-translational processing of preproghrelin appears to produce at least 2 secreted hormones [Kojima et al. 1999, Zhang et al. 2005; comment in Garg A. 2007]: 1. the mature octanoylated ghrelin (acyl ghrelin), derived from residues 24-51 of preproghrelin and acylated on Ser 3 by ghrelin O-acyltransferase (GOAT) [Kojima et al. 1999, Zhu et al. 2006, Ozawa et al. 2007, Yang et al. 2008, Hougland J. 2019], and 2. obestatin, a C-terminally amidated peptide derived from residues 76-98 of preproghrelin [Zhang et al. 2005] that may be subject to carboxypeptidase E and C-terminal peptidyl alpha-amidating enzyme activities [Ozawa et al. 2007]. Solution-state NMR structural dynamics show that ghrelin association with the growth hormone secretagogue receptor (GHS-R) is dependent on the hormone’s N-terminal octanoyl moiety and N-terminal hydrophobic core [Ferre et al. 2019].

A shared axis for ghrelin and insulin signaling in control of glucose metabolism has been put forward [Mani et al. 2019]. Prior studies have shown a role for acyl ghrelin in control of insulin secretion and glucose metabolism [Broglio et al. 2001, Broglio et al. 2004]. In animal models, ghrelin inhibits insulin secretion [Reimer et al. 2003, Dezaki et al. 2004, Salehi et al. 2004, Qader et al. 2005]. In humans, epidemiologic studies have revealed an inverse correlation between ghrelin levels and insulin resistance [Tschöp et al 2001]. Obesity and type 1 diabetes are associated with decreased circulating ghrelin levels [Soriano-Guillén et al. 2004, Martos Moreno et al. 2006], while a single dose of intravenous ghrelin has been shown to increase plasma glucose levels followed by reduction in fasting glucose levels [Broglio et al. 2001], potentially suggesting inhibition of insulin secretion [Müller et al. 2015].

Yet, the metabolic fate of ghrelin remains to be exhaustively characterized. The M13 zinc protease neprilysin (NEP) has been postulated to play a role in ghrelin cleavage on the basis of peptide cleavage products observed in rat serological analysis [De Vriese, 2004 Endocrinology]. However, ghrelin hydrolysis by NEP has not been demonstrated in vivo or in vitro. It remains to be determined whether a significant fraction of ghrelin is retained in circulation following association with the signaling receptor.

A possible role for insulin-degrading enzyme (IDE) in ghrelin proteolysis has not been previously demonstrated. IDE is a member of the M16 subfamily of zinc endopeptidases [Barrett and Rawlings, 2004]. M16 peptidases bear an invariant *HXXEH* zinc-binding and catalytic motif and are termed inverzincins on the basis of motif sequence inversion relative to the *HEXXH* consensus motif of M4 (thermolysin) family proteases [Becker and Roth 1992, Barrett and Rawlings, 2004].

Like other M16 peptidases IDE exhibits broad substrate specificity. Established IDE substrates include insulin [Mirsky et al. 1949], glucagon [Baskin et al. 1974], epidermal growth factor and transforming growth factor alpha [Gehm and Rosner, 1991], and other hormones with roles in homeostasis, cell growth, and metabolism [Becker and Roth 1995]. Common structural features of IDE substrates include a capacity for peptides to fit within the enzyme’s central binding cavity (M_r_ < ~10 kDa), a tendency to adopt beta strand or beta-turn conformations, and hydrophobic or amyloidogenic character proximal to the site of peptide hydrolysis [Shen et al. 2006, Kurochkin IV, 1998, Stefanidis et al. 2018].

Notably, the budding yeast IDE homologs Axl1p and Ste23p support N-terminal cleavage of **a**-factor mating pheromone [Adames et al. 1995]. Yeast **a**-factor is a farnesylated dodecapeptide that is subject to multiple post-translational modifications including N-terminal serine acylation and processive proteolytic cleavage prior to sec-independent export [Michaelis and Herskowitz, 1988; Powers and Michaelis 1986, Michaelis and Barrowman, 2012]. When heterologously expressed in *S. cerevisiae*, mammalian IDE [Kim et al. 2005], the *E. coli* M16A homolog pitrilysin [Alper et al. 2006], and the M16C family *A. thaliana* presequence peptidase PreP [Phillips JE, 2007] all support **a**-factor cleavage. These observations suggest that a capacity to adopt N-terminally acyl lipidated substrates is a specificity factor conserved among homologs of the M16 subfamily.

Here we demonstrate that recombinant human IDE can degrade synthetic octanoylated human ghrelin in vitro. In comparison with prior work, the observed rate of IDE-mediated ghrelin degradation was comparable or superior to the rate of IDE-mediated degradation of insulin-FITC and recombinant amyloid beta 1-40 peptides [Stefanidis et al. 2018]. Moreover, ghrelin degradation appears at least somewhat specific to M16 peptidases such as IDE, as the catalytically active M13 protease NEP did not exhibit ghrelin proteolysis in this study. Ghrelin degradation by IDE is an iterative process. IDE-mediated ghrelin cleavage sites were identified using HPLC and MALDI-TOF mass spectrometry. IDE-mediated ghrelin cleavage sites informed construction of a novel fluorescence-resonance energy transfer (FRET) substrate (Mca-QRVQQRKESKK(Dnp)-OH; Mca: 7-methoxycoumarin; Dnp: 2,4-dinitrophenyl). We anticipate that this substrate may serve as a useful proxy for native ghrelin in future studies of its proteolysis by IDE or other peptidases.

## Materials and methods

### Expression and purification of recombinant enzymes

Isolation of recombinant human IDE, catalytically inactive IDE E111Q, and glutathione S-transferase (GST) were carried out as previously described [Stefanidis et al. 2018]. Amino acid residues M_42_ through L_1016_ of the annotated IDE translation products were expressed in bacterial host BL21 DE3 as C-terminal polyhistidine tagged fusion partner (i.e., IDE_M42-L1016_-6xHis). The IDE E111Q mutant was derived from this construct through targeted mutagenesis of the IDE_M42-L1016_-6xHis clone, achieved by inverse PCR [Stefanidis et al. 2018]. Construction of the C-terminally polyhistidine tagged *S. japonicum* GST (i.e., GST-6xHis) has been previously reported [Considine et al. 2017].

Following transformation with the appropriate expression vector, bacterial cultures were grown to high cell density (0.5-1.0 OD_600_; 37 °C × 195 RPM) and induced for protein expression. Isopropyl-beta-d-thiogalactopyranoside (IPTG) was added to 0.1 μM, cell culture temperature shifted to 25 °C, and cells harvested following overnight protein induction. Bacterial cell pellets were reconstituted in a lysis/wash buffer compatible with immobilized metal affinity chromatography and lysed by ultrasonic disruption. Insoluble material was cleared from the lysate by centrifugation (3200 g x 20 min. at 4 °C). Polyhistidine tagged IDE was isolated from the soluble lysate fraction by immobilized nickel affinity chromatography (NI-IDA resin, Takara Clontech Inc.) in accordance with manufacturer’s instructions. Approximately 100 column volumes of lysis/wash buffer were applied following column loading. Buffers were refrigerated or kept on ice throughout the purification procedure. Purified proteins were eluted with 300 mM imidazole, 300 mM NaCl, and 20 mM sodium phosphate pH 7.4, and held on ice while protein concentration was determined using the BioRad Protein Assay [Bradford MM, 1976]. Elution products were diluted to 50% w/v glycerol and stored at −20 °C.

### Commercial sourcing of synthetic human ghrelin and recombinant human neprilysin

Octanoylated synthetic human ghrelin was purchased from New England Peptide, Inc. (Gardner, MA, USA). Upon receipt, peptide was reconstituted in dimethylsulfoxide (DMSO) to 20 mg/mL, aliquoted and stored at −20 °C. Recombinant human neprilysin was a gift of ACROBiosystems, Inc. (Newark, DE, USA). Upon receipt as a lyophilized powder, neprilysin was reconstituted to 0.2 mg/mL by addition of water and stored at −80 °C.

### SDS-PAGE assay for ghrelin proteolysis

All proteolytic assays in this study were performed in 50 mM Tris(hydroxymethyl)aminomethane (Tris), pH 7.3 and conducted at 37 °C. Conditions for analysis of IDE-mediated ghrelin proteolysis were similar to those used to evaluate IDE-mediated proteolysis of FITC-labelled insulin and amyloid beta peptides [Stefanidis et al. 2018]. Ghrelin degradation assays were performed after separately assembling pre-mixtures containing 75 ng/μL IDE, 75 ng/μL IDE E111Q, 50 ng/μL neprilysin or 50 ng/μL GST, and 2 μg/μL substrate (octanoylated ghrelin). Enzyme and substrate pre-mixtures were preincubated at assay temperature of 37 °C. The assay was initiated by mixing equal parts enzyme and substrate. Proteolytic reactions were quenched at endpoint by addition of SDS-PAGE sample buffer containing 160 mM Tris, pH 6.8, 100 mM EDTA, 32% w/v glycerol, 1.25% w/v β-mercaptoethanol and 0.003% w/v bromophenol blue. Where present for purposes of enzyme inhibition, EDTA, an established IDE inhibitor [Kirschner and Goldberg 1983], was added to the enzyme pre-mixture during assembly. Samples were separated by SDS-PAGE (BioRad Any kD^™^ Mini-PROTEAN^®^ TGX gel, manufacturer’s Tris-glycine buffer system) immediately after the proteolytic assay. Gels were stained using colloidal Coomassie Brilliant Blue G250 [Candiano et al. 2004], and densitometric analysis was performed using a BioRad Gel Doc imaging station and BioRad Image Lab Version 6 graphical analysis software.

### Identification of IDE-mediated ghrelin cleavage sites

Proteolytic assays were performed in the manner above and quenched at endpoint in accordance with established methods [Stefanidis, 2018]. Ghrelin cleavage product separation by HPLC and analysis by MALDI-ToF, samples was achieved as described below. 45 μL of 50 mM Tris, pH 7.3 were added to the quenched proteolysis experiment and samples were injected into the HPLC (Shimadzu, Prominence). Proteolysis products were separated on a Phenomenex Kinetex C-18 analytical column (100 mm x 4.6 mm) using a 1 mL/min flow rate and a linear gradient of 20 to 60 % acetonitrile over 20 minutes. Ghrelin and proteolysis products with elution times between minutes 11 and 14 were collected. The integrated area under the curve for ghrelin and the three proteolysis products was averaged across three runs and plotted for each timepoint. Cleavage sites were confirmed by matrix-assisted laser desorption ionization time-of-flight (MALDI-TOF) mass spectrometry (Shimadzu, Axima Confidence) using α-cyano-4-hydroxycinnamic acid (CHCA) matrix in reflectron mode.

### Mca-QRVQQRKESKK(Dnp)-OH peptide synthesis

A ghrelin derivative FRET substrate was developed using similar design principles as previous insulin B chain and amyloid beta derivative peptide FRET substrates (Stefanidis et al, 2018) that included the major IDE-mediated ghrelin cleavage sites. The FRET substrate Mca-QRVQQRKESKK(Dnp)-OH was synthesized on a PS3 peptide synthesizer (Gryos Protein Technologies) on an Fmoc-Lys(Dnp) Wang resin. After the N-terminal glutamine was added, the final Fmoc was removed, and the resin was washed with DMF and DCM and transferred to a fritted syringe (Torviq). 7-methoxycoumarin-3-carboxylic acid was added at the N-terminus using standard 2-(1H-benzotriazol-1-yl)-1,1,3,3-tetramethyluronium hexafluorophosphate (HBTU) coupling protocols. The FRET peptide was cleaved from the resin with a 5 mL TFA/triisopropylsilane/water (95%/2,5%/2.5%) cocktail for two hours at room temperature. The crude peptide product was precipitated in cold diethyl ether, pelleted by centrifugation, and dried in vacuo. The peptide was HPLC purified using C-18 semi-preparative column (XBridge BEH, Waters) with a flow rate of 3 mL/min over a linear gradient of 20% – 60% B over 20 minutes (Solvent A: 0.1% TFA in H2O; Solvent B: 100% acetonitrile) and monitored at 222 nm. The fractions containing the pure peptide were confirmed by MALDI-TOF using CHCA matrix, then combined and lyophilized.

### FRET assays

Mca-QRVQQRKESKK(Dnp)-OH (FRET ghrelin) proteolysis assays were performed essentially as previously described [Stefanidis, 2018], and adapted for use with the peptide developed for this study. In brief, enzyme (2 ng/μL) and substrate (40 μM) pre-mixtures were separately assembled in 50 mM Tris, pH 7.3 and warmed to 37 °C. The proteolytic reaction was initiated by mixing equal parts enzyme and substrate pre-mixtures (25 μL each). Where indicated, EDTA was added to the enzyme pre-mixture. Upon initiation of the proteolytic experiment, fluorescence was monitored in real time using a BioTek Synergy HT plate reader (excitation 325 ±25 nm, emission 420 ±25 nm, gain setting 90). Half-area black polystyrene 96 well microplates (Corning Costar #3686; Corning Inc., Corning, NY, USA) were used in this study.

Substrate V proteolysis assays were performed as above, alternatively using 20 μM FRET substrate in the pre-mixture. Substrate V (Mca-RPPGFSAFK-Dnp; RND systems, Minneapolis, MN, USA) is a quenched-fluorogenic peptide derivative of bradykinin. Use of this peptide has been previously reported for studies of endothelin converting enzyme (ECE-1), IDE, NEP, and other proteases [Johnson et al., 2000, Sharma et al. 2015, and others].

To identify IDE proteolytic cleavage sites in FRET ghrelin, 50 μL of a quenching solution containing 500 mM EDTA and 2% TFA were added to the 50 μL enzymatic assay mixture 10 minutes into the proteolysis experiment. Samples were stored at −80 °C prior to analysis of peptide cleavage sites by MALDI-TOF mass spectrometry. A 25 μL aliquot of the quenched reaction mixture was prepared for MALDI-TOF mass spectrometry with a C18 reverse-phase ZipTip (Millipore) following the manufacturer’s protocols. MALDI-TOF was performed using a CHCA MALDI matrix in reflectron mode.

### Circular dichroism analysis of IDE substrates

To investigate the feasibility of using circular dichroism (CD) to obtain kinetic parameters for the IDE-dependent degradation of acyl ghrelin (Ivancic et al., 2018), CD spectra of the substrate in 50 mM Tris pH 7.3 in the absence or presence of 30% 2,2,2-trifluoroethanol (TFE), a solvent that enhances helix formation in short peptides with intrinsic alpha-helical propensity (Lazo and Downing, 1997), were recorded. To test the effect of TFE on IDE’s activity, CD spectra of insulin in the presence of IDE at a substrate:enzyme ratio of 100:1 were also acquired. All spectra were recorded at 37 °C on a JASCO J-815 spectropolarimeter using quartz cuvettes with a path length of 1 mm. Acquisition parameters include wavelength range of 260 to 200 nm and 4 scans per spectrum with each scan recorded with a 1 nm bandwidth and a 4 s averaging time.

## Results

Synthetic octanoylated human ghrelin was degraded by recombinant human insulin-degrading enzyme (IDE; Fig. 1). However, ghrelin was not proteolyzed by IDE bearing the inactivating glutamate 111 to glutamine (E111Q) catalytic site point mutation, neprilysin (NEP), or glutathione S-transferase (GST). The molecular weight of ghrelin is ~3400 kDa. Consistent with peptide degradation by a zinc-dependent metallopeptidase, disappearance of a low MW fragment in the presence of catalytically active IDE was sensitive to and limited by pretreatment of the enzymatic activity with ethylenediaminetetraacetate (EDTA), a metal chelating agent and established IDE inhibitor [Kirschner and Goldberg 1983].

**Figure 1.**
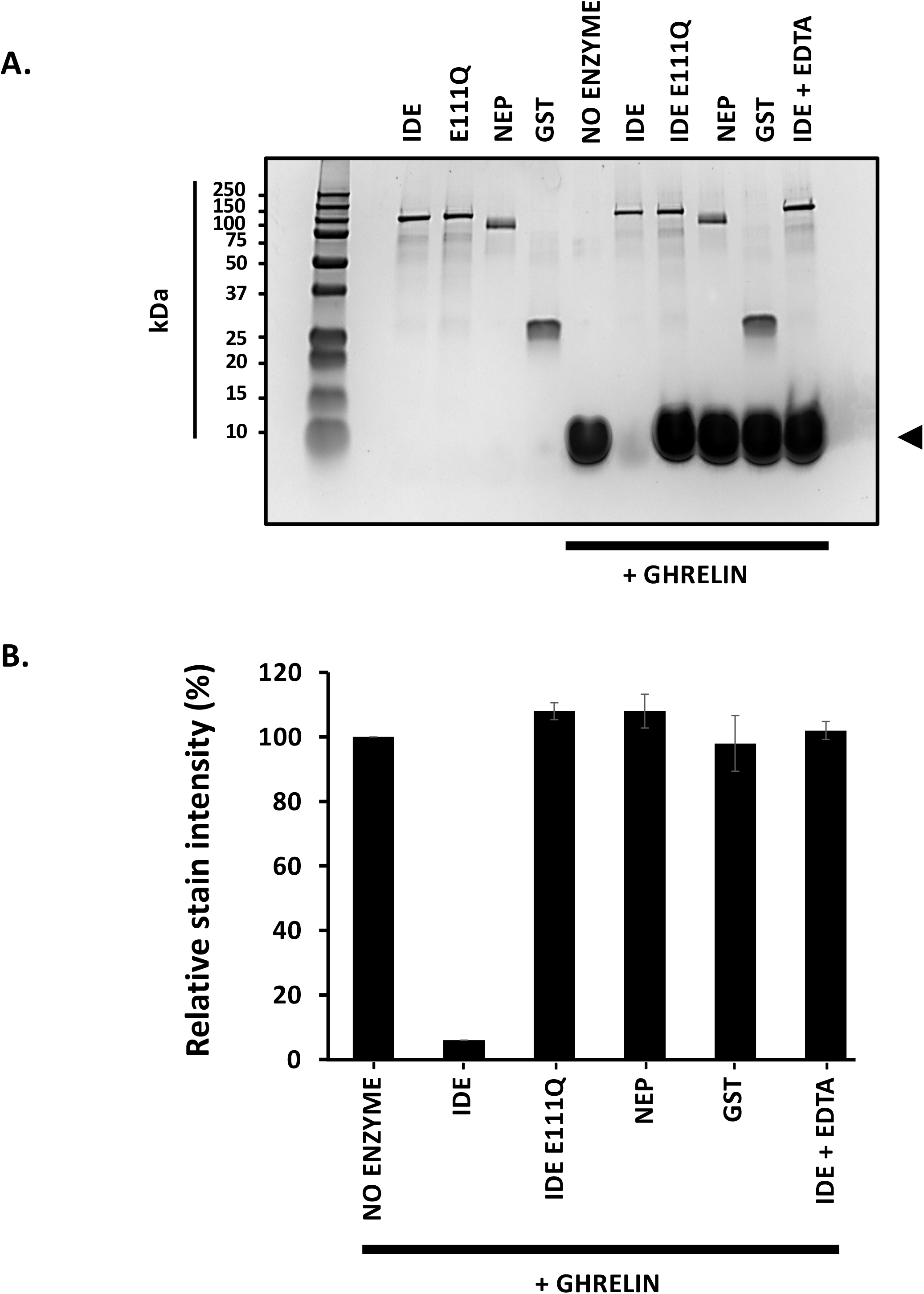
Ghrelin proteolysis by recombinant insulin-degrading enzyme (IDE). **A.**SDS-PAGE analysis of synthetic acyl ghrelin degradation following 1 hr coincubation with IDE, a catalytically inactive IDE mutant (IDE E111Q), neprilysin (NEP), glutathione S-transferase (GST), or IDE pretreated with ethylenediaminetetraacetate (IDE+EDTA) as a metal chelating agent and metalloprotease inhibitor. The migratory position of ghrelin prior to digestion is indicated with a black arrowhead (◄) Gel load amounts correspond with input of 10 μg octanoylated synthetic ghrelin and 0.38 μg IDE or IDE E111Q or 0.25 ug NEP or GST to the initial assay mixture. The gel was stained with colloidal Coomassie Brilliant Blue G-250. Molecular weights of intact, full-length peptides and proteins are as follows: octanoylated ghrelin, 3371 Da; IDE and IDE E111Q, 110 kDa; NEP, 86 kDa; GST, 26 kDa. **B.** Densitometric analysis inclusive data presented in Fig. 1A was performed using BioRad Image Lab Version 6 densitometry software (2 experimental replicates).

Ghrelin degradation progresses towards completion over time in the presence of IDE, but not in the presence of the catalytically inactive IDE E111Q mutant or NEP (Fig. 2A), consistent with enzyme-mediated proteolysis by active IDE only. Densitometric analysis (Fig. 2B) suggests the specific activity of ghrelin proteolysis by IDE is 140 ± 7.4 (standard deviation) nmol min^-1^ mg^-1^ across the first 5 minutes of the experimental time course. These values reflect the approximate rate of peptide degradation as judged by disappearance of stain intensity. However, they do not account for the rate of individual cleavage events within the multistep, iterative process of IDE mediated peptide degradation.

**Figure 2.**
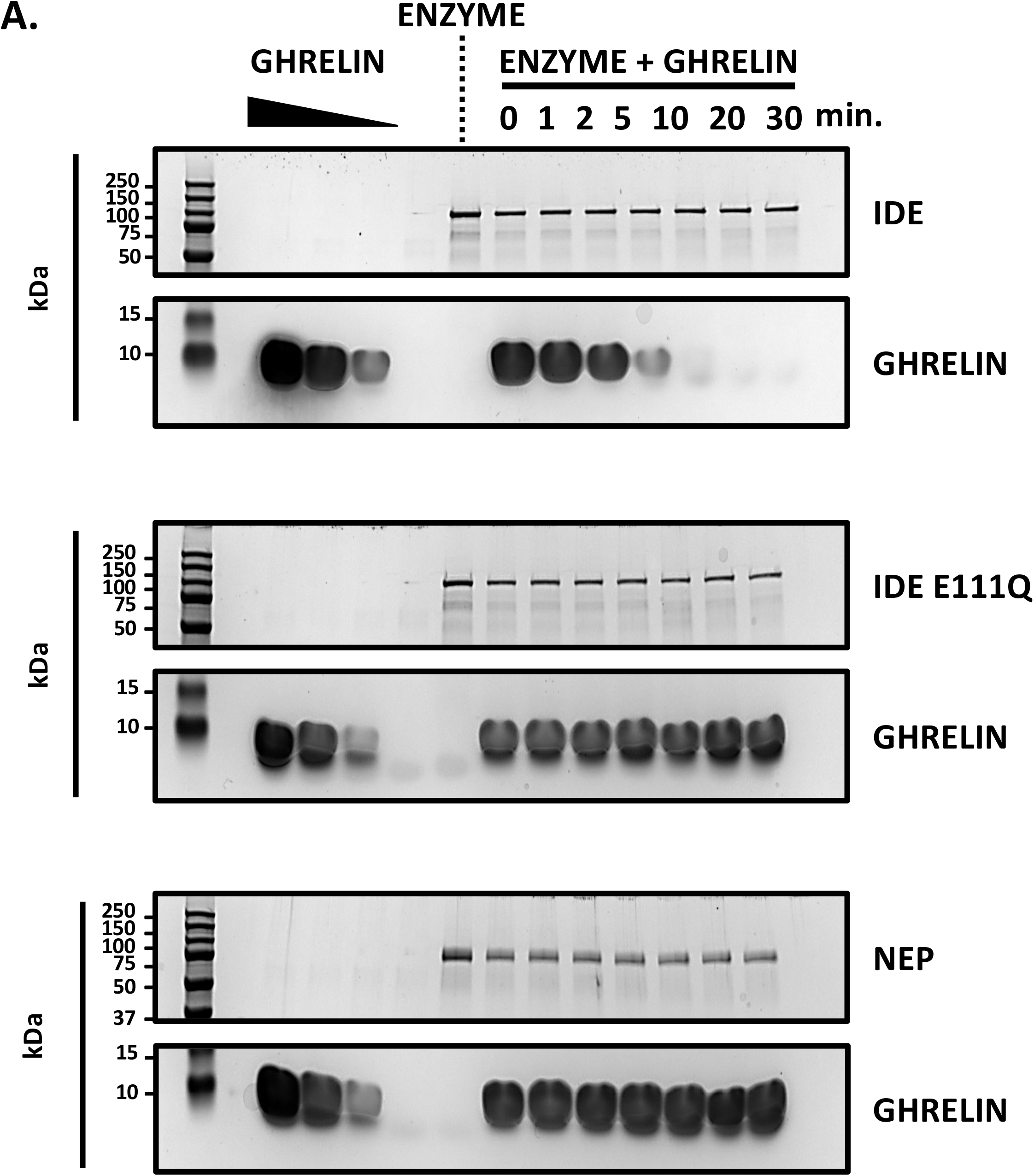
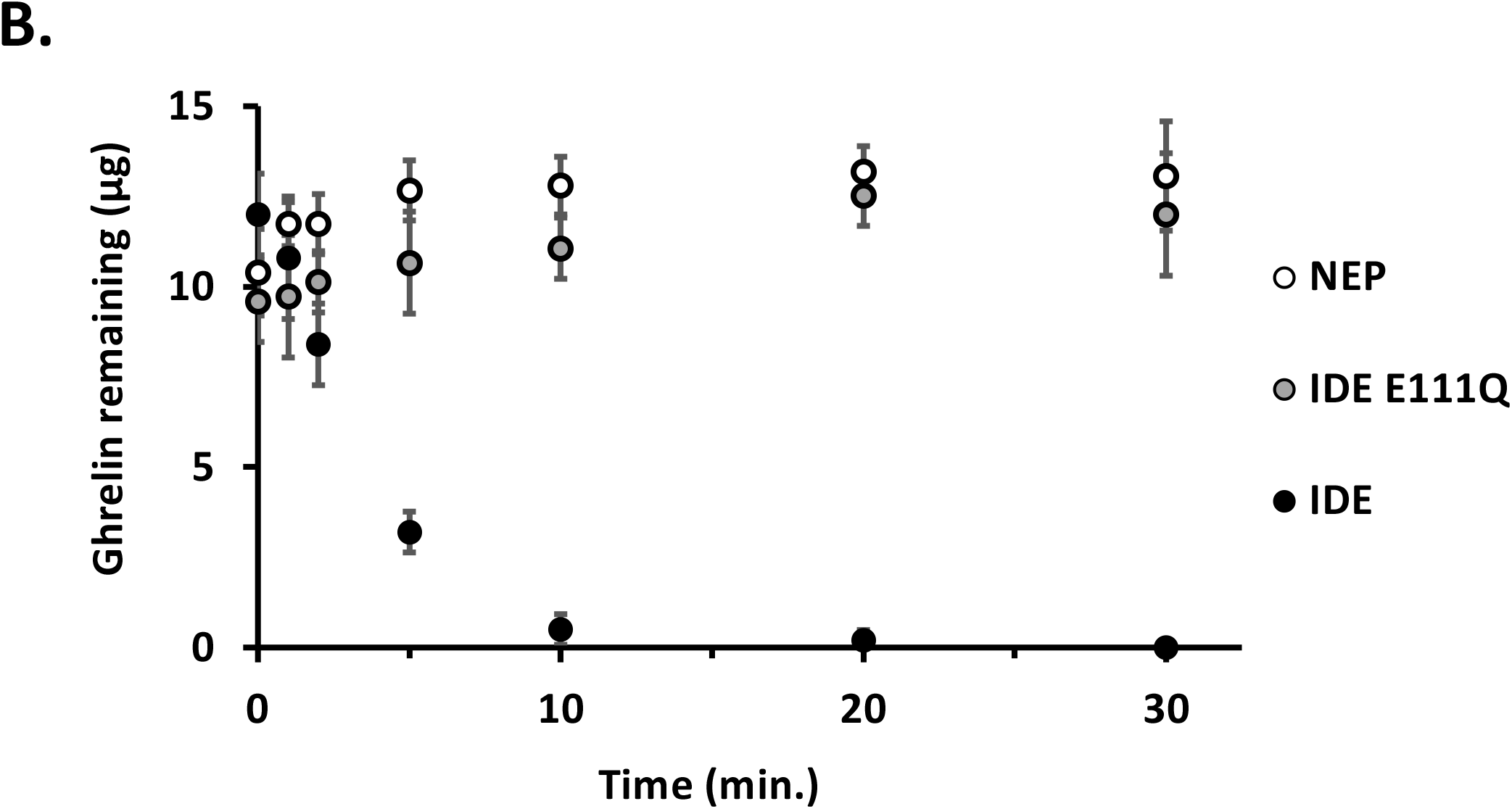
Kinetics of ghrelin degradation by IDE. **A.** SDS-PAGE analysis of ghrelin proteolysis following coincubation with IDE, IDE-E111Q catalytic mutant, and neprilysin (NEP). Reactions were terminated at the timepoint indicated. **B.** Densitometric analysis inclusive data shown in Fig. 2A was performed by peak integration using BioRad Image Lab Version 6 densitometry software (≥ 2 experimental replicates). The rate of ghrelin hydrolysis by IDE from 0 to 5 minutes was 140 ± 7.4 (standard deviation) nmol min^-1^ mg^-1^.

To examine the order and location of ghrelin cleavage sites targeted by IDE, peptide hydrolysis was evaluated by HPLC and MALDI-TOF (Fig. 3). Ghrelin proteolysis by IDE appears processive, with initial cleavage sites between residues Q13/ Q14, Q14/R15 and possibly Q10/R11 preceding later cleavage events (chromatograms in Fig. 3A and 3B). The proteolysis products resulting from cleavage between Q14/R15 and Q13/Q14 increase rapidly in the first 20 minutes of the proteolytic assays and then plateau or decrease for the Q13/Q14 and Q14/R15 sites, respectively; however, there is a lag in appearance of the third cleavage site between Q10/R11 (Fig. 3C). Further analysis is required to confirm whether the Q10/R11 proteolytic product is produced after initial Q14/R15 and Q13/Q14 cleavage. Mass spectra for proteolysis products are shown in Fig. 3D-F.

**Figure 3.**
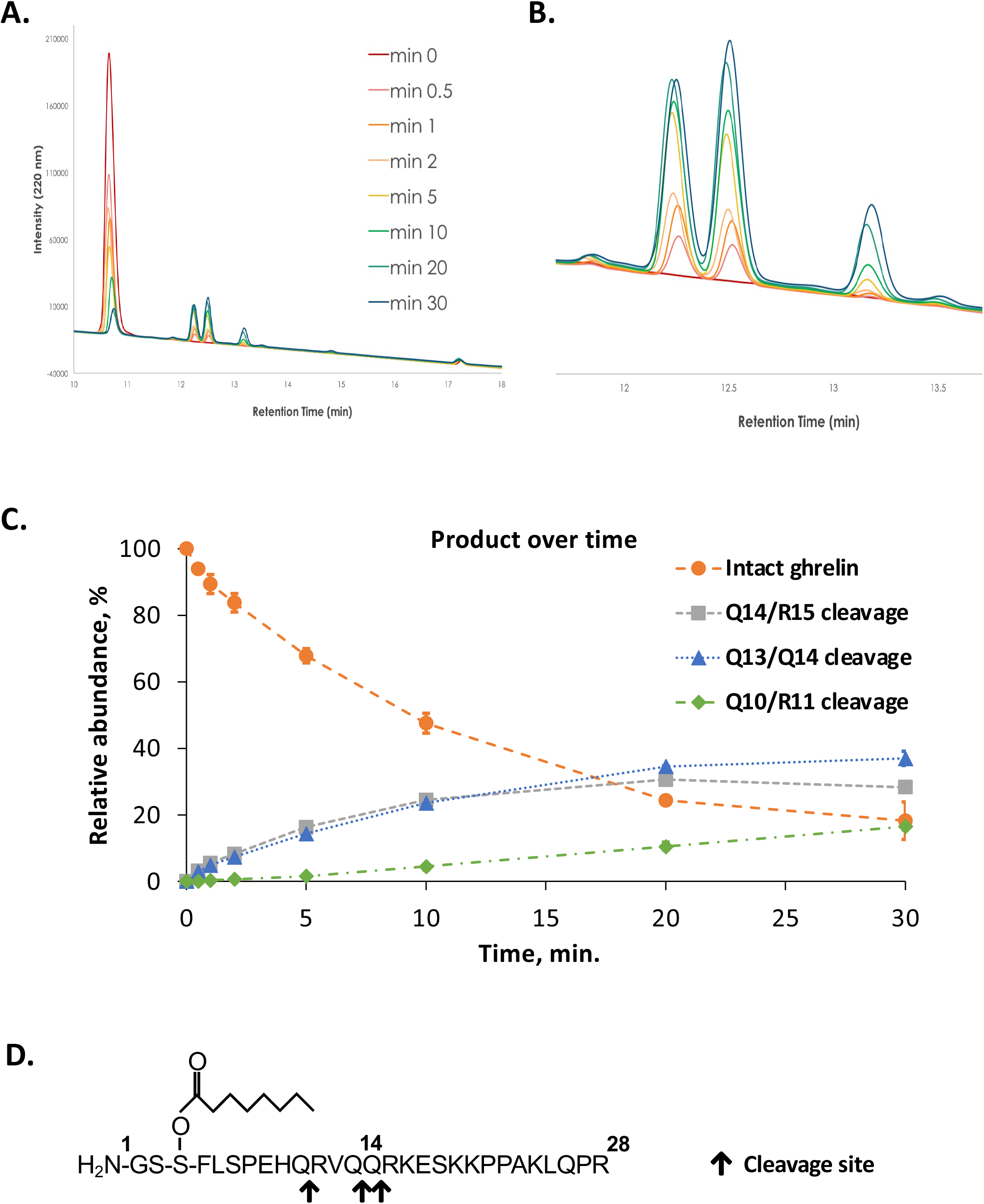
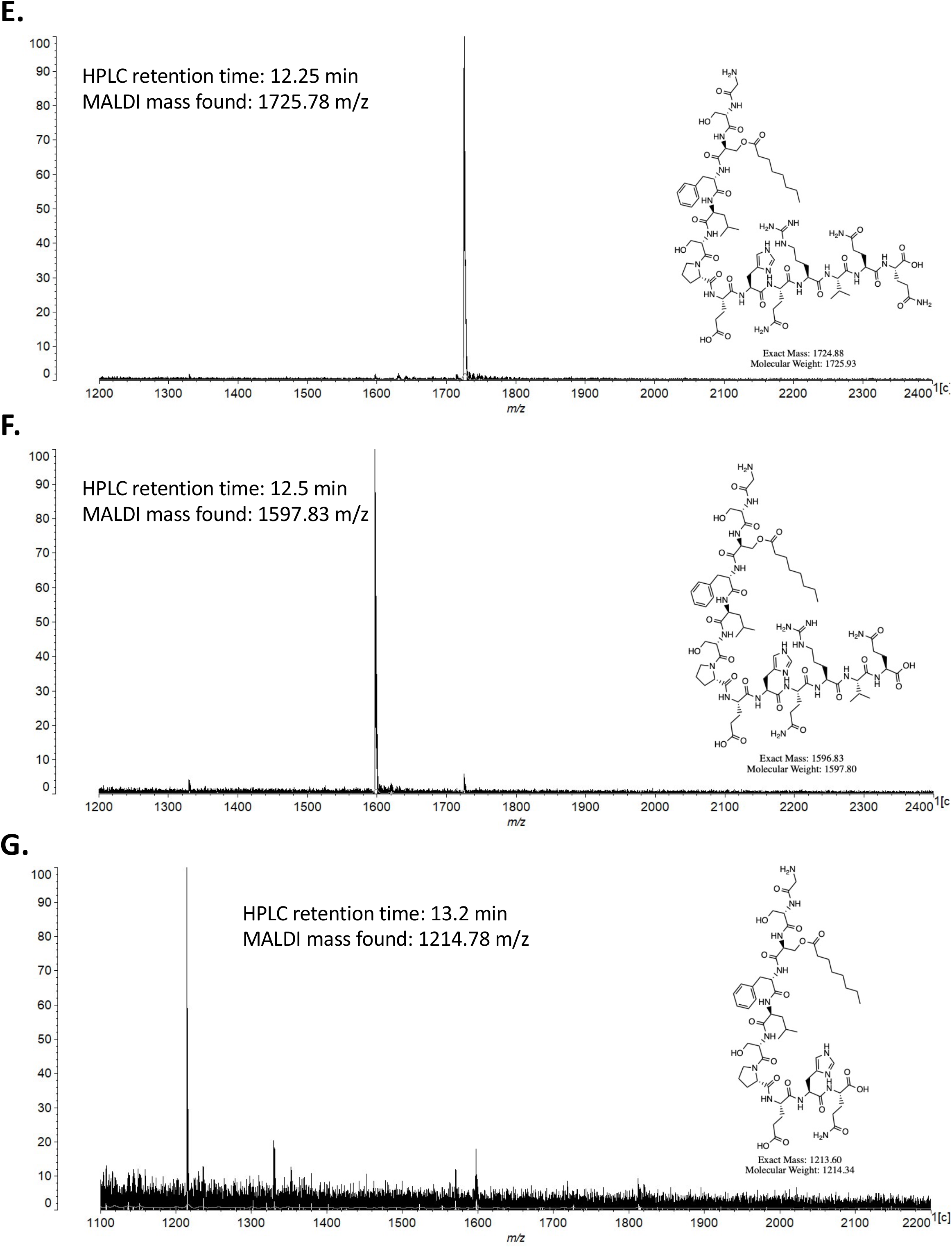
IDE-mediated ghrelin cleavage sites. **A.** HPLC chromatograms revealing IDE-mediated ghrelin cleavage over time. Chromatographic traces for each timepoint are shown. **B.** Inset from chromatogram shown in 3A, revealing peak elution profiles from ~11.5-14 min. Peaks corresponding with single-site cleavage at Q14/R15, Q13/Q14, and Q10/R11 are predominant. **C.** Ghrelin cleavage products over time. Relative quantities of intact peptide and cleavage products are inferred from peak integration of data above. **D.** Schematic representation of IDE-mediated ghrelin cleavage sites. Predominant early cleavage sites are indicated with black arrows (**↓**). **E.** Identification of ghrelin cleavage fragment with HPLC retention time of 12.5 min. (m/z 1725.78) by MALDI-TOF. **F.** Identification of ghrelin cleavage fragment with retention time of 12.5 min. (m/z 1597.83). **G.** Identification of ghrelin cleavage fragment with retention time of 13.2 min (m/z 1214.78).

Experimentally determined proteolytic cleavage sites informed construction of the Mca-QRVQQRKESKK(Dnp)-OH internally quenched fluorogenic peptide (FRET ghrelin; Fig. 4). FRET ghrelin contains amino acid residues immediately N- and C-terminal to the predominant IDE-mediated ghrelin cleavage sites at Q13/Q14 and Q14/R15 (Fig. 4A; grey arrows). Mca-QRVQQRKESKK(Dnp)-OH synthesis was performed using standard solid-phase peptide chemistry, purified by HPLC (Fig. 4B), and verified by MALDI-TOF (Fig. 4C). FRET ghrelin proteolysis by IDE was evaluated by fluorogenic assay (Fig. 4D). As with the full length acylated hormone, FRET ghrelin proteolysis by IDE was inhibited by addition EDTA to the enzyme pre-mixture. Mca-QRVQQRKESKK(Dnp)-OH peptide cleavage sites were similar to those observed within acyl ghrelin, occurring at positions 5analogous to residues Q13/Q14 and Q14/R15 of the native hormone (Fig. 4E). Assay standardization is shown (Fig. 4F). FRET ghrelin was cleaved more rapidly by IDE than by NEP (Fig. 4C), despite NEP exhibiting superior proteolytic activity towards the bradykinin derivative FRET peptide Substrate V (Mca-RPPGSFAFK-Dnp; Fig. 4G).

**Figure 4.**
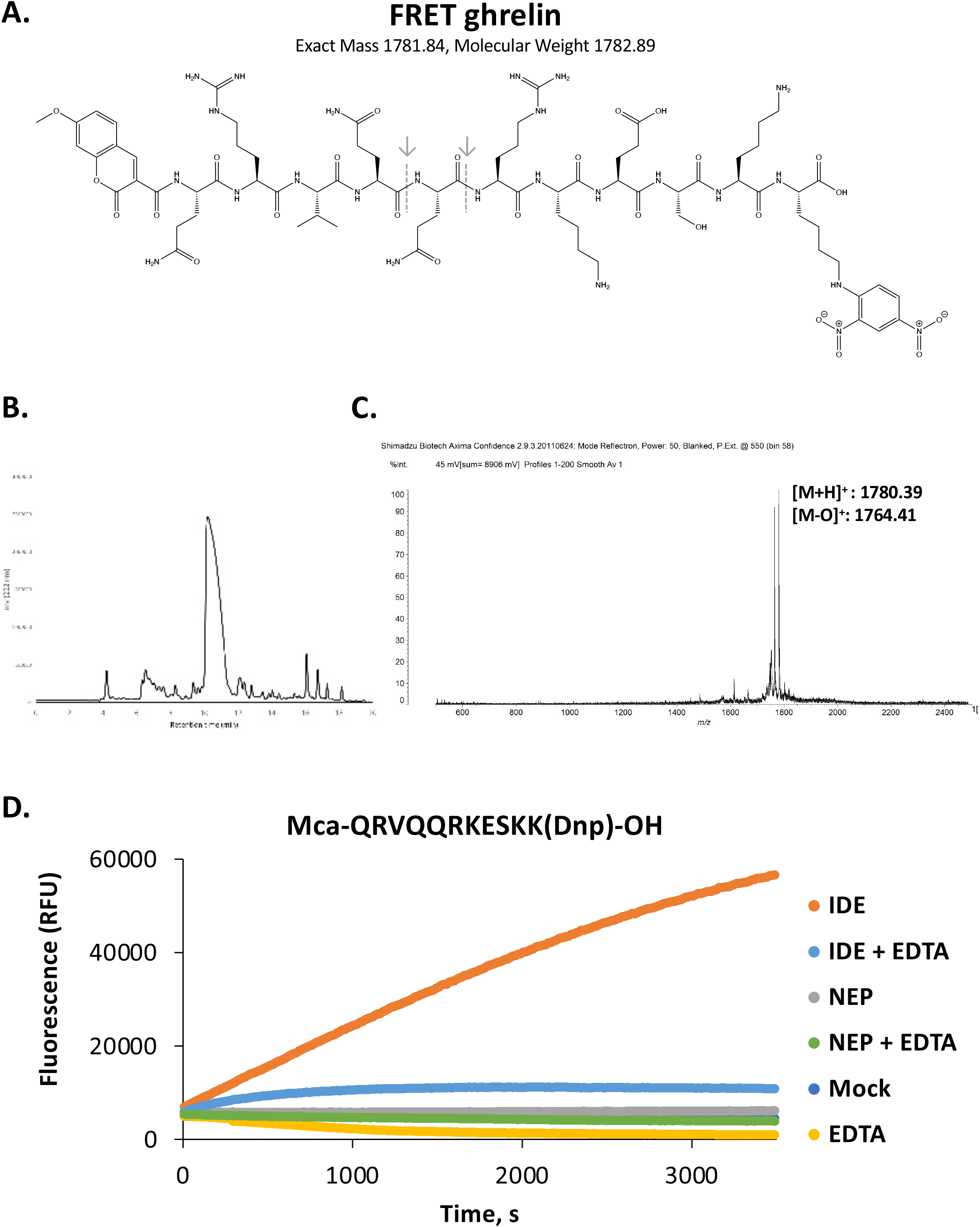
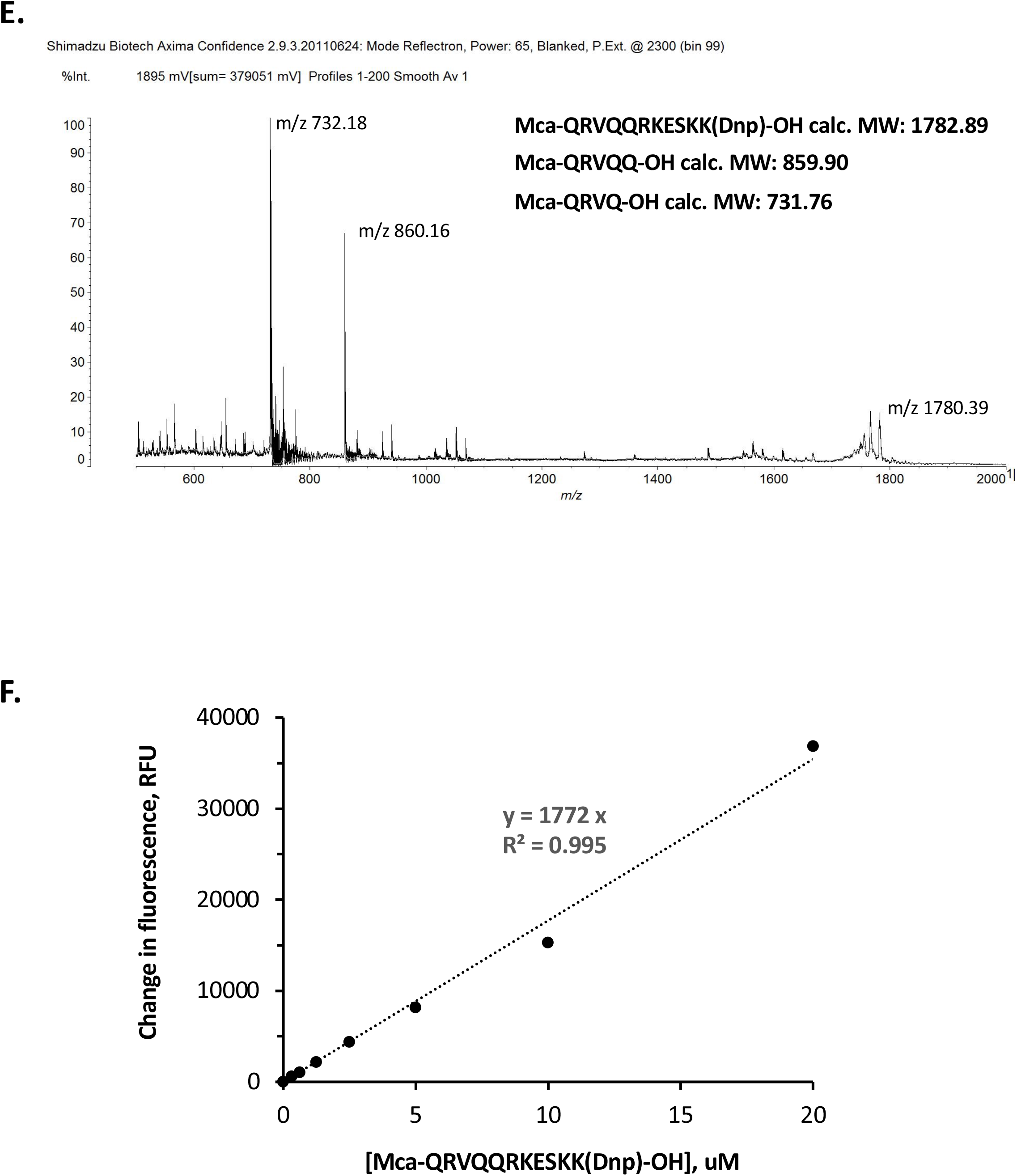
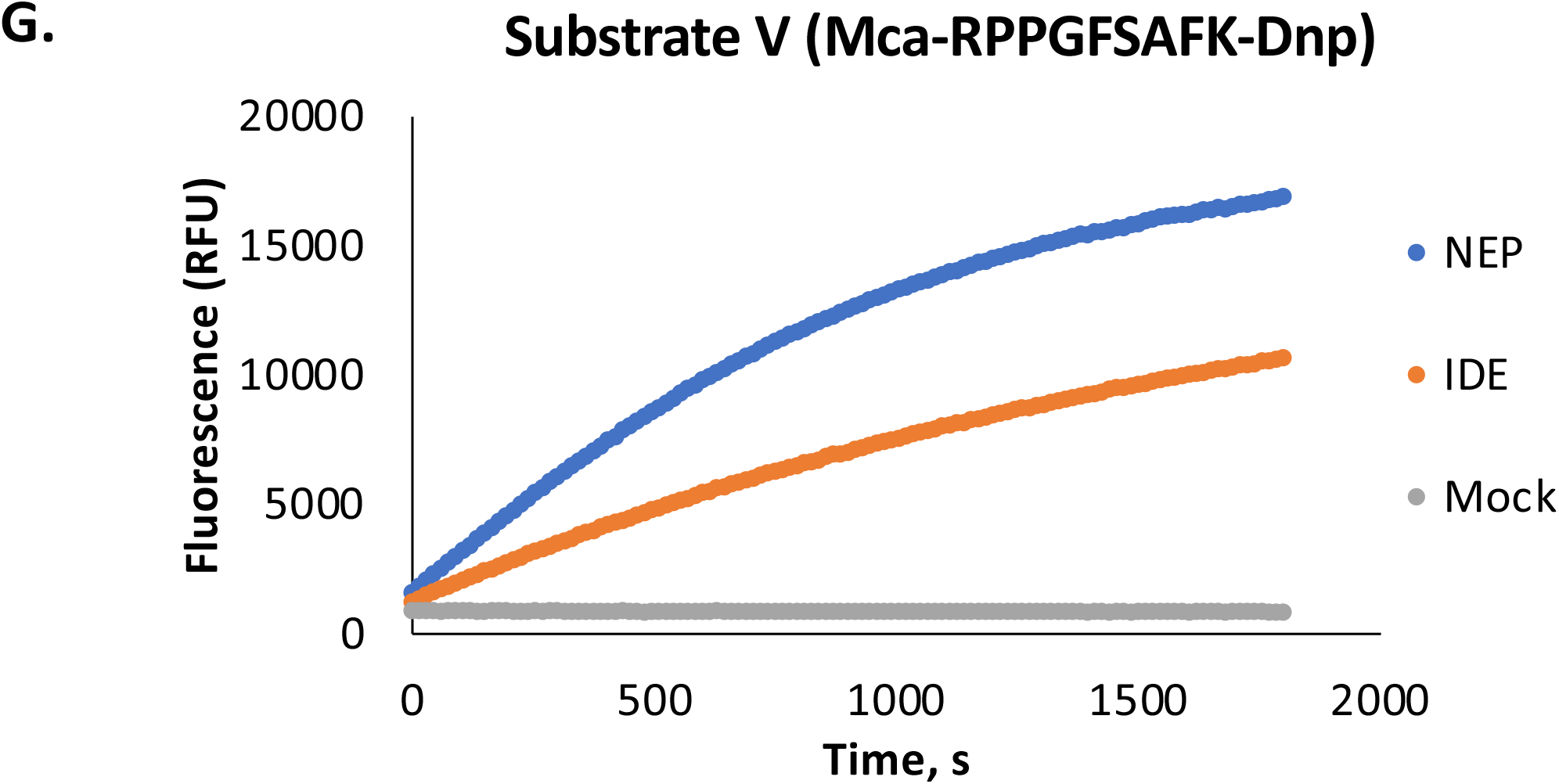
Design, synthesis and proteolysis of a ghrelin derivative quenched-fluorogenic peptide. **A.** Representation of the Mca-QRVQQRKESKK(Dnp)-OH peptide (FRET ghrelin; Mca: 7-methoxycoumarin-3-carboxylic acid, Dnp: 2,4-dinitrophenyl) made for and used in this study. Positions of IDE mediated cut sites are indicated with dashed lines and grey arrows (**↓**). **B.** Reverse-phase (C18) HPLC chromatogram for isolation of the Mca-QRVQQRKESKK(Dnp)-OH peptide following solid-phase synthesis and cleavage. **C.** Confirmation of Mca-QRVQQRKESKK(Dnp)-OH synthesis and purification by MALDI-TOF. **D.** Kinetic analysis of Mca-QRVQQRKESKK(Dnp)-OH cleavage by real time fluorescence assay (excitation: 325 ± 25 nm, emission: 420 ± 25 nm). The Mca-QRVQQRKESKK(Dnp)-OH peptide alone (mock; deep blue) does not exhibit significant change in fluorescence over the time course, and is largely overlapped by the NEP, and NEP+EDTA datasets. **E.** Identification of FRET ghrelin cleavage fragments by MALDI-TOF following proteolysis of Mca-QRVQQRKESKK(Dnp)-OH by IDE. Major cleavage fragments Mca-QRVQQ-OH (m/z 860) and Mca-QRVQ-OH (m/z 732) are identified. The full-length FRET peptide (m/z ~1780) remains detectable following incomplete hydrolysis. **F.** Assay standardization relating FRET ghrelin fluorescence at proteolysis assay endpoint as a function of input substrate concentration. **G.** Proteolysis of bradykinin derivative commercial Substrate V (Mca-RPPGFSAFK-Dnp) confirms catalytic activity of both IDE and NEP.

IDE exhibited inferred half-occupancy by Mca-QRVQQRKESKK(Dnp)-OH at peptide concentration of 7.49 μM (K_half_; 95% CI 5.91-10.8 μM, Fig. 5). Specific activity of proteolysis was 34.6 nmol min^-1^ mg^-1^ (95% CI 30.5-41.5 nmol min^-1^ mg^-1^). These values may be compared with those reported towards insulin-derivative (Mca-VEALYLVCGEK(Dnp)-OH, K_half_:1.26 ± 0.14 μM, specific activity: 116 ± 6 nmol min^-1^ mg^-1^) or amyloid beta-derivative (Mca-QKLVFFAEDVK(Dnp)-OH, K_half_: 3.16 ± 0.69 μM 183±12.6 nmol min^-1^ mg^-1^) FRET substrates under similar conditions of analysis [Stefanidis et al., 2018].

**Figure 5.**
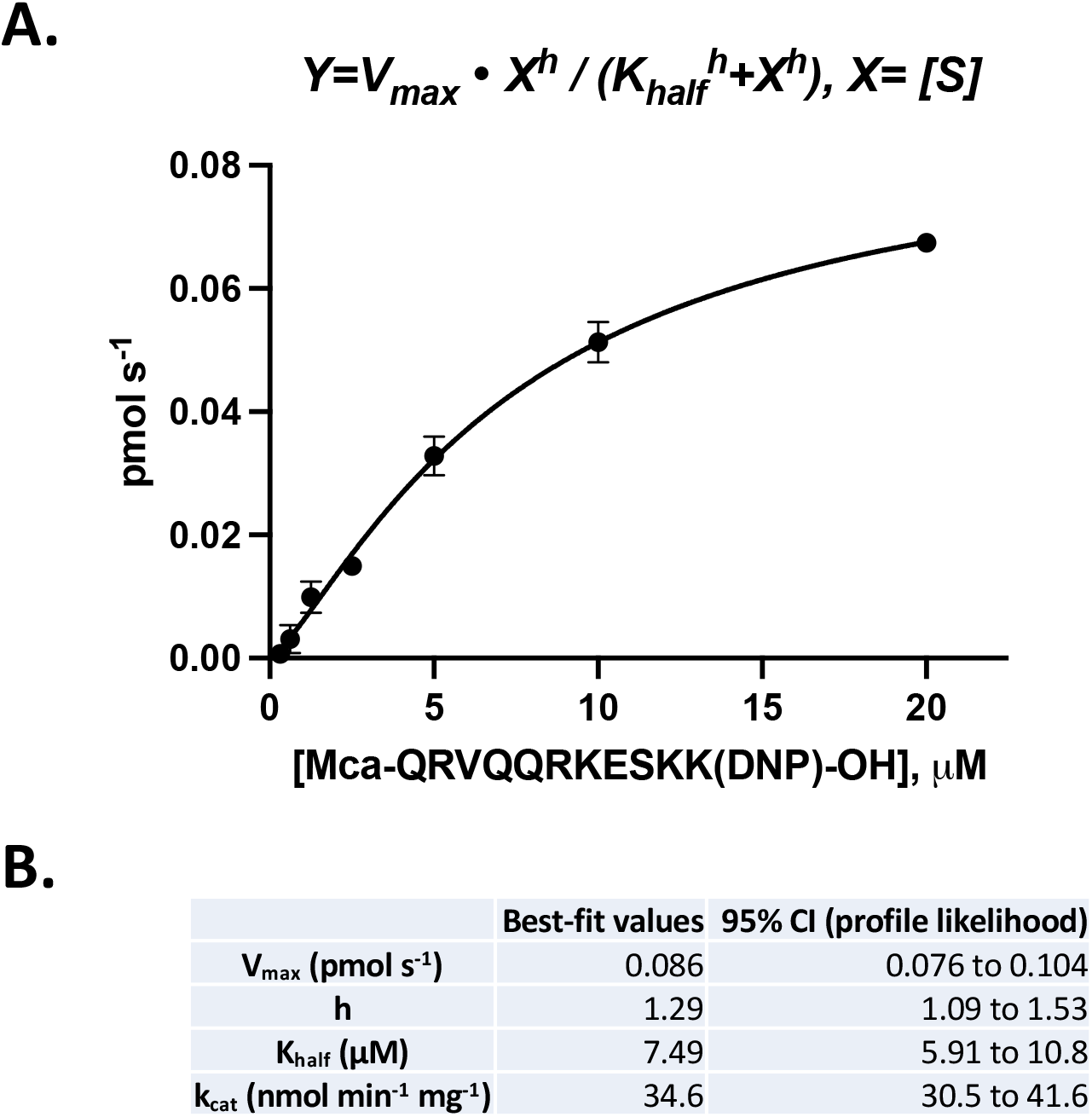
Kinetic parameters of FRET-ghrelin proteolysis by IDE. **A.** Saturation curve for IDE-mediated proteolysis of Mca-QRVQQRKESKK(Dnp)-OH. **B.** Kinetic parameters of proteolysis. Reported values were determined using the nonlinear regression function of GraphPad PRISM^™^.

Efforts to investigate the kinetics of ghrelin proteolysis by IDE using circular dichroism were unsuccessful (Figure 6). Ghrelin bears significant alpha-helical in 30% TFE (Fig. 6A) but appears largely unstructured in the 50 mM Tris pH 7.3 proteolysis buffer used throughout this study (Fig. 6B). IDE does not degrade insulin in 30% TFE (Fig. 6C), despite retaining significant insulin degradation activity in 50 mM Tris pH 7.3 proteolysis buffer as indicated by the loss of helical dichroic signal at 222 nm (Fig. 6D; see also Ivancic et al., 2018). Liquid chromatography (LC) analysis of the insulin + IDE proteolysis experiment performed in 30% TFE did not provide evidence of peaks corresponding with proteolytic cleavage fragments, consistent with inactivity of IDE in 30% TFE solvent (not shown). Further work may therefore be warranted to adapt CD for investigation of ghrelin proteolysis.

**Figure 6.**
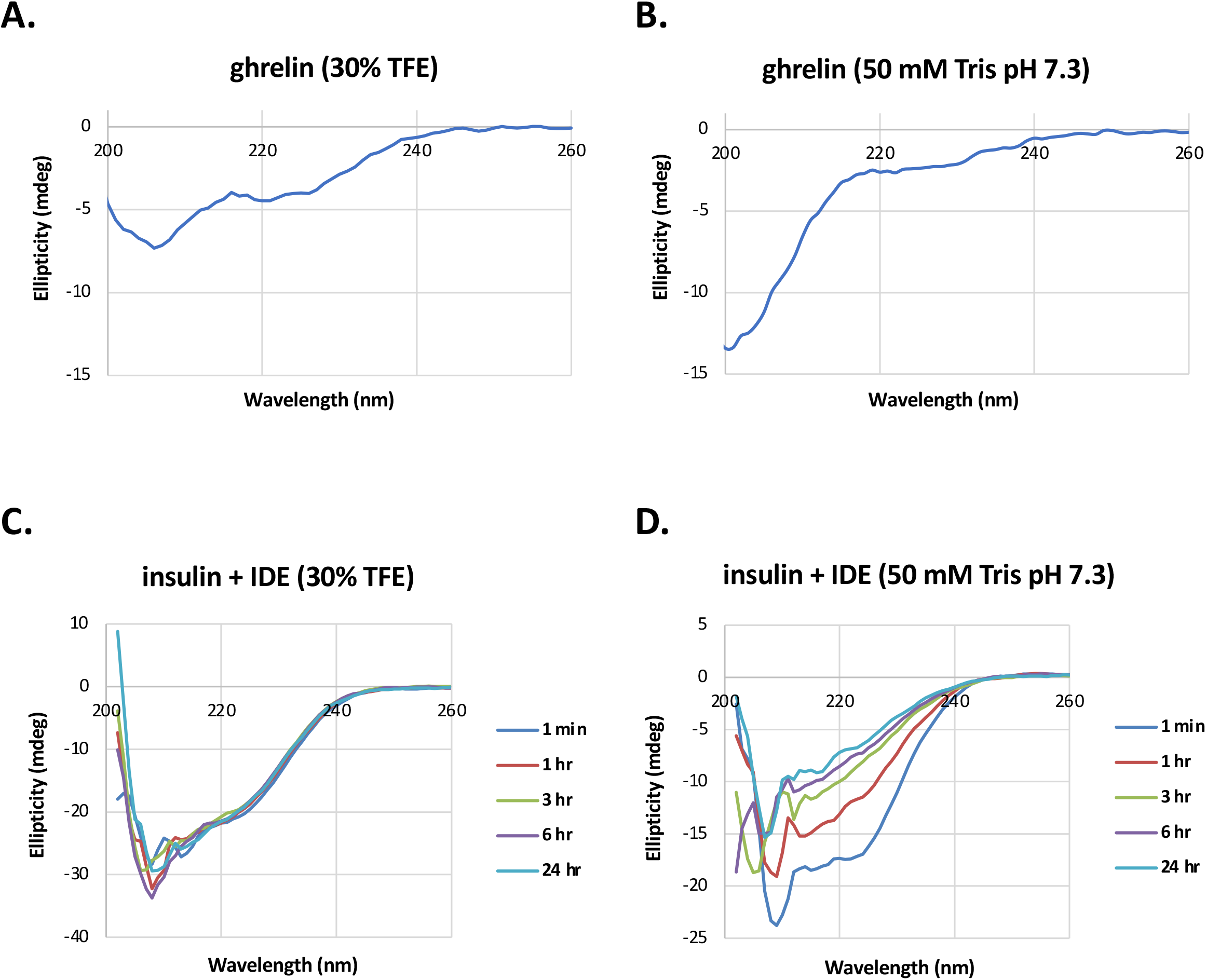
Circular dichroism structural analysis of IDE substrates. **A.** Circular dichroism (CD) spectrum of acyl ghrelin in 30% TFE (2,2,2-trifluoroethanol). **B.** Acyl ghrelin in 50 mM Tris, pH 7.3. **C.** Insulin and insulin degrading enzyme (IDE) in 30% TFE. Substrate to enzyme ratio is 100:1. **D.** Insulin and IDE in 50 mM Tris, pH 7.3. Substrate to enzyme ratio is 100:1. The intensity of insulin’s helical dichroic signal at 222 nm decreases with digestion time. All CD spectra were recorded at 37 °C.

## Discussion

This study demonstrates ghrelin proteolysis by insulin-degrading enzyme (IDE). This study expands the in vitro specificity of human IDE to include N-terminally acyl-lipidated peptides with hydrophobic amino acid moieties. Consistent with a role for IDE as an important regulator of cell energy homeostasis, demonstration of the capacity for ghrelin proteolysis by IDE adds to an expanding group of peptide substrates that includes insulin [Mirsky et al. 1949], glucagon [Baskin et al. 1974], epidermal growth factor and transforming growth factor alpha [Gehm and Rosner, 1991], and others. The physiological significance, if any, of IDE-mediated ghrelin proteolysis remains to be demonstrated.

## Acknowledgements

This work was supported by funding from Sacred Heart University College of Arts and Sciences, Department of Chemistry and Physics to BJA and DDB. This work was also supported by National Science Foundation Grant CHE-1624774 to JES-C, for which BJA served as contributor, and by a National Institutes of Health Grant R15AG055043 to NDL. Recombinant human neprilysin was a gift from ACROBiosystems, Inc. (Newark, DE, USA). BJA thanks James L. Hougland (Syracuse University, Syracuse, NY) for correspondence and scientific insight, and Walter K. Schmidt, Jr. (University of Georgia, Athens, GA) for continuing correspondence and mentorship.

